# Moderate Endurance Exercise Increases Arrhythmia Susceptibility and modulates Cardiac Structure and Function in a Sexually Dimorphic manner

**DOI:** 10.1101/2023.08.21.554195

**Authors:** Sharon A George, Katy Anne Trampel, Kelsey Brunner, Igor R Efimov

## Abstract

**Background:** While moderate endurance exercise has been reported to improve cardiovascular health, its effects on cardiac structure and function are not fully characterized, especially with respect to sexual dimorphism. We aimed to assess the effects of moderate endurance exercise on cardiac physiology in male versus female mice.

**Methods:** C57BL/6J mice of both sexes were run on a treadmill for six weeks. ECG and echocardiography were performed every two weeks. After six weeks of exercise, mice were euthanized, and triple parametric optical mapping was performed on Langendorff perfused hearts to assess cardiac electrophysiology. Arrhythmia inducibility was tested by programmed electrical stimulation. Left ventricular (LV) tissue was fixed, and RNA sequencing was performed to determine exercise-induced transcriptional changes.

**Results:** Exercise-induced LV dilatation was observed in female mice alone, as evidenced by increased LV diameter and reduced LV wall thickness. Increased cardiac output was also observed in female exercised mice but not males. Optical mapping revealed further sexual dimorphism in exercise-induced modulation of cardiac electrophysiology. In female mice, exercise prolonged action potential duration and reduced voltage-calcium influx delay. In male mice, exercise reduced the calcium decay constant, suggesting faster calcium reuptake. Exercise increased arrhythmia inducibility in both male and female mice, however, arrhythmia duration was increased only in females. Lastly, exercise-induced transcriptional changes were sex-dependent: females and males exhibited the most significant changes in contractile versus metabolism-related genes, respectively.

**Conclusions:** Our data suggest that moderate endurance exercise can significantly alter multiple aspects of cardiac physiology in a sex-dependent manner. While some of these effects are beneficial, like improved cardiac mechanical function, others are potentially pro-arrhythmic.

## INTRODUCTION

Cardiovascular disease (CVD) is the leading cause of death in the United States, accounting for ∼875,000 deaths annually.(1) A sedentary lifestyle is among many risk factors for heart disease. Thus, researchers and doctors continue to recommend exercise as a primary approach to the prevention and management of CVDs. Several studies have reported the cardiac response to exercise in healthy (athletes) and diseased hearts.(2–5) During exercise, the faster rate of muscle contraction increases metabolic demand. This is supported by an increased blood supply of nutrients and oxygen to the muscles, including the heart muscle. To provide the increased blood supply, the heart undergoes dramatic remodeling, including left ventricle (LV) dilatation and increased heart rate. Some of these effects of exercise cause chronic cardiac remodeling that persists even during resting periods, such as dilatation, hypertrophy, inflammation, altered cardiac mechanics, electrical remodeling, and altered SA and AV nodal function.(6–8) The cardiac response to exercise is highly dependent on the type, intensity, and duration of the exercise, and diet.

Different types of exercise include endurance, strength, balance, and flexibility. They can be further classified as low, moderate, and high-intensity exercise.(9) It has been recommended that a greater amount of physical activity can significantly reduce the risk of disease through the body’s ability to increase aerobic capacity and improvement of muscle strength and function. However, high intensity exercise causes electrical remodeling in the hearts of athletes’ resulting in bradycardia, atrioventricular block, T wave inversion in electrocardiogram (ECG) and ectopy. (10) On the other hand, low and moderate endurance exercise lowers blood pressure and cholesterol, reduces risk of Type 2 Diabetes and some types of cancer, and is associated with increased life span (11,12). The American Heart Association also details health benefits associated with mental health and cognition; more specifically, one’s memory and attention span can improve with increased exercise, along with fewer symptoms of depression and anxiety. While the positive overall effects of moderate exercise on health are well-established, its effects on cardiac structure and electrical and mechanical function are unknown.

Age, sex, and genetic predisposition can influence the physiological response to exercise.(13,14) However, sex differences in cardiac structure and function response to moderate intensity exercise are mostly unknown. Male-only studies in biological research vastly outnumber female-only studies at a ratio of about 4 to 1.(15) Even when studies include both sexes, less than half have incorporated sex into their analysis. Recent studies have identified sex differences in multiple aspects of cardiac physiology, including electrophysiology, metabolism, and mechanics.(16–19) Women have longer QT intervals, smaller ejection fraction (EF), and are more dependent on fatty acids metabolism for ATP production compared to males, in addition to smaller heart size and numerous proteomic and transcriptional differences. (17,20,21) Many of these effects are attributed to sex hormones – estrogen and testosterone. These sex differences are further amplified by CVDs or stress. Sex differences in athletes’ heart have been discussed in a review by St Pierre, (20) where in response to high intensity exercise, male and female hearts undergo significant remodeling, some of which were observed in male hearts alone, some in female hearts alone and others in both males and females. Here, we hypothesized that moderate intensity exercise also causes significant remodeling of cardiac structure and electrical/mechanical functions in a sex-dependent manner.

This study investigated differences in three aspects of the cardiac response to moderate-level endurance exercise in male and female mice: structural, mechanical, and electrical. Treadmill running was chosen for the endurance exercise regimen and cardiac physiology was assessed by echocardiography, electrocardiography, and triple-parametric optical mapping. The combination of these modalities allowed for a thorough characterization of cardiac structural, mechanical, and electrical function which was then correlated with transcriptional changes.

## METHODS

All animal protocols were approved by the Institutional Animal Care and Use Committee at George Washington University and in accordance with the National Institutes of Health Guide for Care and Use of Laboratory Animals.

### Experimental Groups

Male and female wild-type C57BL/6J mice (The JAX Laboratory, 000664) were randomly assigned into four groups: sedentary male (n = 11), sedentary female (n = 11), exercised male (n = 9), and exercised female (n = 9) (Figure 1A). At 15 weeks of age, exercised mice began 4 days of exercise training to determine if all mice could be trained to run on a custom rodent treadmill (Conquer Under Desk Portable Electric Treadmill Walking Pad with custom running lanes built on top). During the training period, mice were run at progressively increasing speeds (up to 1 km/h) and durations (up to 20 mins), and air puffs from a compressed air can were applied to direct the mice to run in the correct direction. At the end of the training period, baseline (Week 0) echocardiogram and ECG recordings were acquired from the exercised mice and age-matched sedentary controls. This was followed by 6 weeks of the exercise regimen where mice ran for 45 minutes per day, 5 days per week, for 6 weeks at a speed of 1 km/hr with no incline. This exercise regimen was chosen because it was previously classified as moderate intensity in mice based on studies that reported ∼75% maximum oxygen consumption (VO_2_max) and maximal lactate steady state accomplished with similar protocols in mice.(22,23)

**Figure 1:**
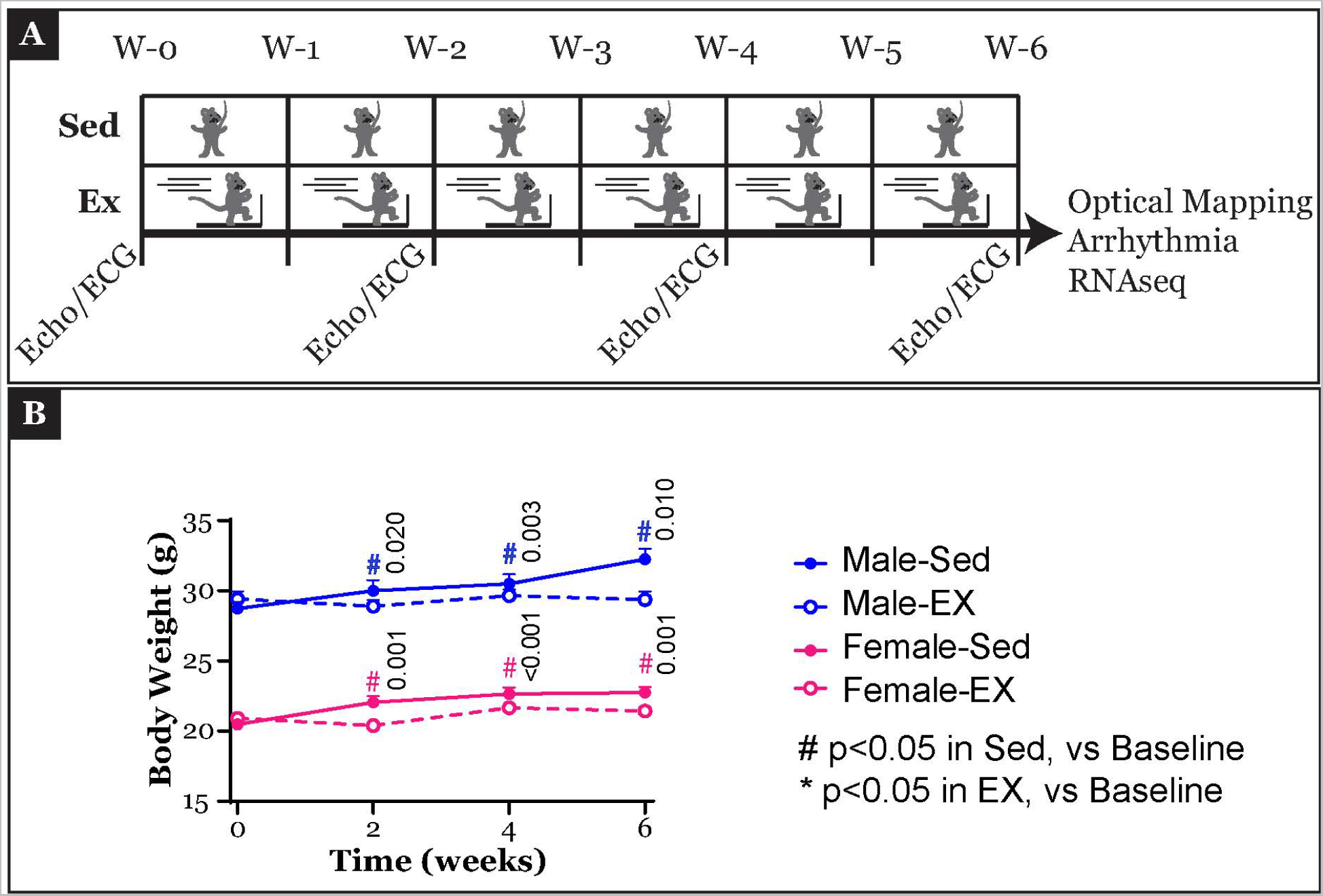
Experimental Protocol. **A)** Different experimental protocols and timelines applied to sedentary (Sed) and exercised (EX) mice over 6 weeks (W0 to W6) are illustrated. **B)** Changes in the total body weight of mice in the four experimental groups were recorded biweekly and are plotted over 6 weeks. Three-way ANOVA tests were performed to determine significant differences in body weight data presented in panel B, with time, sex, and treatment (Sed, EX) as variables. For post-hoc analysis, to compare changes in body weight in each mouse over the duration of the experimental protocol, paired Student’s t-tests were performed, and significant p-values are indicated in the Figure. Benjamini-Hochberg correction was applied for multiple comparisons correction. p<0.05 denotes significance. The sample size for each group is as follows: Male sedentary = 11, Male exercised = 9, Female sedentary = 11, Female exercised = 9.

### Echocardiography

At the end of weeks 0, 2, 4, and 6, exercised and sedentary mice were anesthetized using 2.5% isoflurane at 1 ml/min oxygen flow using an EZ anesthesia machine. Repeated measurements at these time points allowed the determination of time course of exercise-induced modulation of cardiac structure and mechanical function. Mice were weighed to determine total body weight and transferred to an imaging stage heated to 37⁰C with continued isoflurane oxygen mixture flow through a nose cone. Chest hair was removed, and ultrasound gel (Aquasonics, Clear) was applied to the left of the sternum. Transthoracic M-mode echocardiography of the LV was performed using Vevo 3100 (VisualSonics) Ultrasound Machine, and data was analyzed using VevoLAB2.1.0. The following parameters were measured/calculated from the echocardiograms: LV end-systolic diameter (LVESD, mm), LV end-diastolic diameter (LVEDD, mm), LV posterior wall thickness during systole (LVPWs, mm), LV posterior wall thickness during diastole (LVPWd, mm), LV Mass (mg), stroke volume (SV, μl), ejection fraction (EF, %), fractional shortening (FS, %), cardiac output (CO, ml/min), and heart rate (HR).

### Electrocardiography

ECGs were recorded from the anesthetized mice during each echocardiography session at weeks 0, 2, 4, and 6, by attaching the mouse limbs to the four ECG electrodes on the heated platform. The following parameters were measured from the ECG using a custom Matlab program: P wave duration (P, ms), P-R interval (PR, ms), QRS duration (QRS, ms), Q-T interval (QT, ms), and R-R interval (RR, ms). Definitions for each parameter measured from mouse ECG were previously described.(24) QT_C_ intervals were not reported because the mouse QT interval has been reported to not change significantly with heart rate.(25)

### Langendorff Perfusion

After the 6-week exercise regimen, male and female exercised (n = 6 each) and sedentary (n = 5 and n = 6, respectively) mice were anesthetized with isoflurane and sacrificed following cervical dislocation. The hearts were excised after thoracotomy, and the aorta was cannulated. Each heart was attached to a Langendorff perfusion system, hung vertically in a tissue chamber, and retrograde perfused with modified Tyrode’s solution (in mM, NaCl 130, NaHCO_3_ 24, NaH_2_PO_4_ 1.2, MgCl_2_ 1, Glucose 5.6, KCl 4, and CaCl_2_ 1.8 at pH 7.4, 37⁰C and bubbled with carbogen – 95% O_2_ and 5% CO_2_). Coronary pressure was maintained at ∼80 mmHg by adjusting the coronary flow rate between 1 and 2 ml/min. ECG electrodes in the tissue chamber recorded pseudo-ECGs. The hearts were paced at 1.5x stimulation threshold and 2 ms duration at 150ms basic cycle lengths (BCL) using a platinum pacing wire placed in the middle of the anterior surface of the heart.

### Triple-Parametric Optical Mapping

The *ex vivo* Langendorff perfused hearts were allowed a 10-minute equilibration period in the tissue chamber. This was followed by sequential perfusion of voltage- and calcium-sensitive dyes. Over a 3–5-minute period, 1 ml of RH237 (30 µl of 1 mg/ml RH237 stock solution + 970 µl Tyrode’s solution, Biotium 61018) was injected into the heart through the dye port. After a 5-minute washout period, the heart was injected with 1 ml of Rhod2-AM (30 µl of 1 mg/ml Rhod2-AM stock solution + 30 µl Pluronic F-127 + 940 µl Tyrode’s solution, Thermo Fischer Scientific R1244 and Biotium 59055, respectively) over a period of 3-5 minutes and followed by a second 5-minute washout period. Baseline optical recordings were acquired. This was followed by 15 µM blebbistatin (Cayman Chemicals 13186) perfusion to uncouple the heart electromechanically. The heart was then illuminated by two light-emitting diode (LED) light sources at 365 nm (Mightex Systems, LCS-0365-04-22) and 520 ± 5 nm (Prizmatix, UHP-Mic-LED-520) wavelength to excite nicotinamide adenine dinucleotide (NADH) autofluorescence and RH237/Rhod2-AM, respectively. Emitted light was captured and directed to three complimentary metal-oxide semiconductor (CMOS) cameras (MiCAM, SciMedia) as previously described.(26)

Custom-developed Rhythm 3.0 was used to analyze NADH, transmembrane potential, and intracellular calcium-related parameters.(26) A total of ten different parameters were measured. From transmembrane potential, rise time (V_m_ RT), action potential duration (APD_80_), transverse and longitudinal conduction velocity (CV_T_ and CV_L_), and anisotropic ratio (AR) were determined. Rise time (Ca^2+^ RT), calcium transient duration (CaTD_80_), calcium decay constant (Ca τ), and V_m_-Ca^2+^ delay were calculated from intracellular calcium recordings. NADH level was calculated as the ratio of NADH intensity at a given BCL to the NADH intensity from the first recording (baseline) from the same heart.

### Arrhythmia Inducibility

Arrhythmia inducibility was assessed by subjecting the Langendorff-perfused hearts to burst pacing while pseudo-ECGs were recorded to characterize arrhythmias. Arrhythmias were distinguished by type and duration. Ventricular tachycardia (VT) was identified as a wide complex and an increased heart rate, greater than 100 beats per minute. Ventricular fibrillation (VF) was identified by ECG waves varying in amplitude and morphology with no identifiable wave parameters. An arrhythmia episode was recorded if its duration was > 5 seconds. Incidences of VT and VF, as well as total arrhythmia duration, were determined from the pseudo-ECG traces. Hearts were also optically mapped during the arrhythmia episodes.

### RNA Sequencing

Tissue from the LV of 3 sedentary and 3 exercised mice of each sex was fixed in RNAlater^TM^ (Millipore Sigma, MFCD03453003), and RNA sequencing was performed by Genewiz from Azenta Life Sciences. Differentially expressed genes (DEGs) were identified using the DESeq2 package, where the significance level was set to less than 0.05. Volcano plots, top ten upregulated genes and pathways were determined for each group. For volcano plots and top upregulated genes, log2 fold change was calculated as Sed value minus EX value (Figure 6) and male value minus female value (Figure 7). Therefore, a positive log2 fold change corresponds to upregulation in Sed or male group while negative log2 fold change corresponds to upregulation in EX or female group. For the upregulated genes and pathways graphs, only the top ten candidates are illustrated in the figures. Additionally, for the top ten genes, non-annotated genes were excluded from the graphs. Upregulated pathways in each group were identified by performing KEGG (Kyoto Encyclopedia of Genes and Genomes) analysis using the clusterProfiler package, where significance level was set to less than 0.05.

### Statistical Analysis

Statistical analyses were performed using Graphpad Prism. For data presented in Figures 1B (body weight), 2B (echocardiography) and 3B (electrocardiography), three-way ANOVA tests with time, sex and treatment (Sed, EX) as the three variables, time as a repeated measure was performed. Results are presented in Supplement 3. On this data set, paired Student’s t-tests were performed as post hoc analysis, to compare values at Weeks 2, 4, and 6 in each mouse to its own baseline value at Week 0. The results of these tests are discussed in the Results section and significant p-values are indicated in the figures. Finally, male versus female comparison, was performed by unpaired, Student’s t-tests and these results are discussed in the Results section. For data presented in Figures 4C (triple-parametric optical mapping) and 5B (arrhythmias), two-way ANOVA test were performed, with sex and treatment as the variables, Time was not a variable in these data sets as these measurements required explantation of the heart and was only performed at the end of the 6-week experimental protocol. For these data sets, unpaired Student’s t-tests were performed to detect statistically significant differences in control versus sedentary and male versus female groups. The former results are discussed in the figures while the latter in the Results section. Benjamini-Hochberg correction was applied for multiple comparison corrections. P<0.05 denotes significance. Data are represented as mean ± standard deviation unless stated otherwise.

For RNAsequencing data, DESeq2 package was used to perform analysis which uses Wald test to determine p-values and applies Benjamini-Hochberg correction to determine p-adj values. For the volcano plots, top genes and pathways plots, p-adj cut off value was < 0.05. Log2 fold change cut off value of 2 was used for top genes and o for volacano and KEGG plots.

## RESULTS

Mouse body weight was tracked at baseline (Week 0) and during the 6-week exercise period. While the body weight of exercised male and female mice remained steady during this period, sedentary male and female mice demonstrated a progressive increase in body weight with age (Figure 1B). Additionally, all male mice had higher body weight compared to females in both the sedentary (p-values = <0.0001 for Weeks 0 to 6) and exercised (p-values = <0.0001 for Weeks 0 to 6) groups.

### Sex-dependent structural remodeling in response to exercise

Transthoracic M-mode echocardiography was performed, and changes in LV structure were characterized (Figure 2A). Echocardiographic parameters were normalized to baseline (Week 0) values and are illustrated in Figure 2B top panel. In exercised female mice, LVEDD (3.4±0.2 vs 3.8±0.2 and 3.7±0.1 mm at weeks 4 and 6, respectively) and LVESD (2.2±0.2 vs 2.5±0.2 and 2.5±0.2 mm at weeks 4 and 6, respectively) were both increased after 4 and 6 weeks of exercise suggesting enlarged ventricles. Furthermore, exercise also decreased LVPWs in these female mice at weeks 2 and 6 (1.2±0.1 vs 1.1±0.1 and 1.1±0.1 mm at weeks 2 and 6, respectively). In male exercised mice, only LVPWd was increased at week 6 (0.7±0.1 vs 0.8±0.1 mm). In sedentary female mice, LV mass increased at week 4 and returned to baseline by week 6. Finally, among exercised mice, LVEDD and LVESD were significantly higher (p = 0.014 and 0.009, respectively) whereas LVPWs was lower (p = 0.011) in females compared to males.

**Figure 2:**
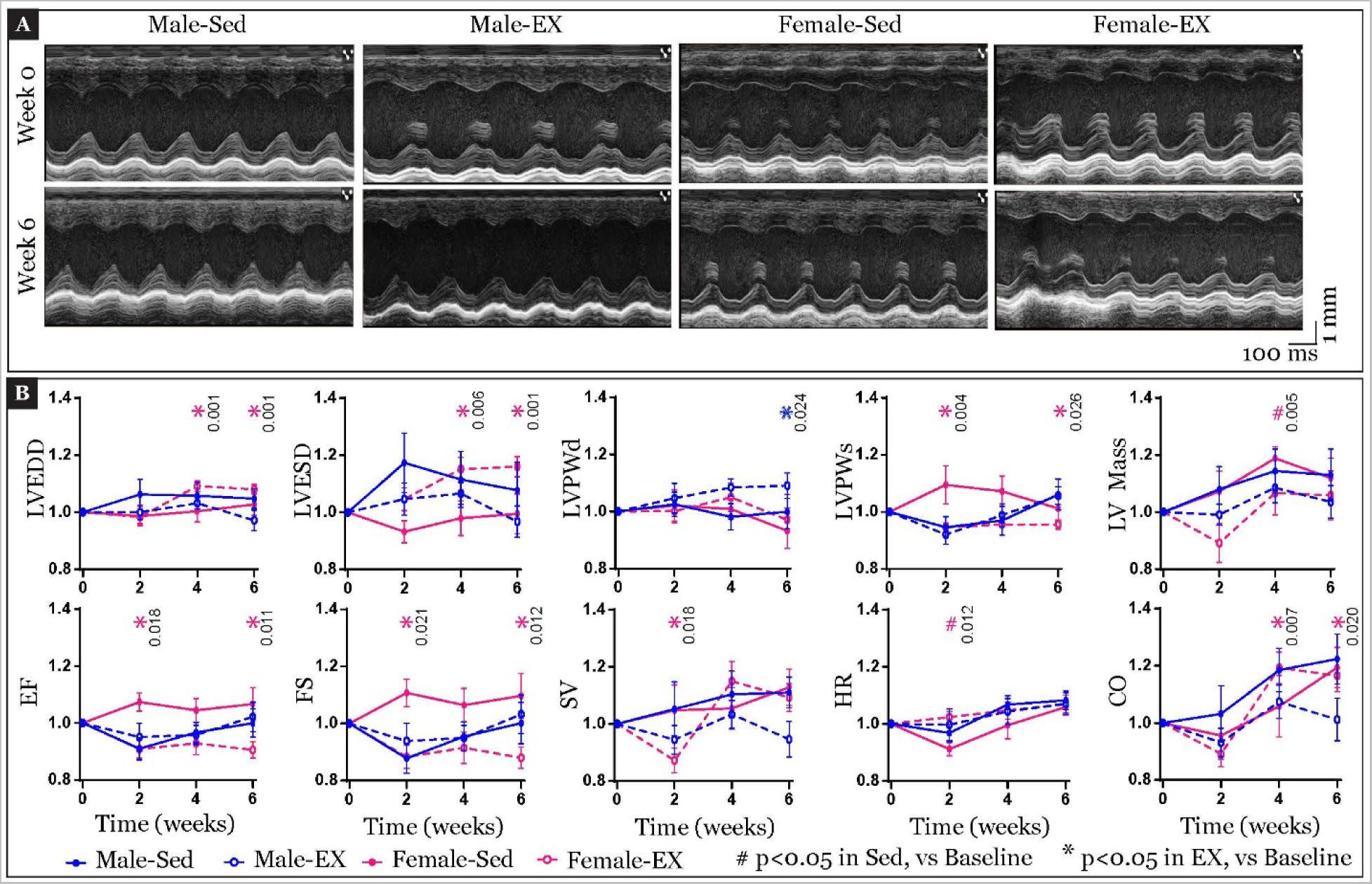
Echocardiographic assessment of cardiac response to exercise. **A)** Echocardiography was performed biweekly and representative echocardiograms from male and female, Sed and EX mice at baseline (Week 0, top panels) and after the 6-week protocol (Week 6, bottom panels) are illustrated. Echocardiograms in each column’s top and bottom panels are from the same mouse. **B)** Summary data of structural (top) and mechanical (bottom) parameters measured from the echocardiograms, normalized to each mouse’s own baseline value prior to start of experimental protocol is plotted over time. LVEDD: left ventricular end diastolic volume, LVESD: left ventricular end systolic volume, LVPWd: left ventricular posterior wall thickness during diastole, LVPWs: left ventricular posterior wall thickness during systole, LV Mass: left ventricular mass, EF: ejection fraction, FS: fractional shortening, SV: stroke volume, HR: heart rate, CO: cardiac output. Three-way ANOVA tests were performed to determine significant differences in echocardiographic data presented in panel B, with time, sex, and treatment as variables. Paired, two-tailed Student’s t-tests were performed for post hoc analysis to determine significant changes induced by experimental protocol in the same mouse over time, and significant p values are indicated in the figure. Benjamini-Hochberg correction was applied for multiple comparisons correction. p<0.05 denotes significance. The sample size for each group is as follows: Male sedentary = 11, Male exercised = 9, Female sedentary = 11, Female exercised = 9.

Taken together, these data suggest that exercise causes LV dilatation in female mice while it could result in LV hypertrophy in male mice. Thus, structural remodeling in exercised mice is sex dependent.

### Sex-dependent mechanical remodeling in response to exercise

Next, exercise-induced changes in cardiac mechanical function were characterized (Figure 2A). Echocardiographic parameters were normalized to baseline (Week 0) values and are illustrated in Figure 2B bottom panel. Significant mechanical remodeling was observed only in female exercised mice. Specifically, EF (66±6 vs 60±9 and 60±6% at weeks 2 and 6, respectively) and FS (36±5 vs 32±7 and 31±4% at weeks 2 and 6, respectively) were reduced in female exercised mice at weeks 2 and 6. While SV was initially reduced at week 2 (33±3 vs 29±4 μl), an increasing trend was observed at week 4 (p = 0.06). This produced a significant increase in CO in these female exercised mice at weeks 4 and 6 (14±2 vs 16±2 and 16±2 ml/min at weeks 4 and 6, respectively). An initial drop in HR was also observed in female sedentary mice at week 2, but this parameter was restored to baseline by week 4. No significant changes were observed in any male mice, exercised or sedentary. Comparing males to females, EF and FS were significantly higher in sedentary females at Week 2, however, this sex differences was not observed in any other parameters.

Reductions in EF observed in female mice do not necessarily indicate reduced mechanical function. But interpreting these results in conjunction with the LV dilatation reported above could indicate that the decrease in EF is an effect of the increased diastolic filling (EF denominator) rather than reduced SV (EF numerator). Regardless, it is of note that, once again, the effects of exercise were sex-dependent and were predominantly observed in female mice.

### Sex-dependent electrical remodeling in response to exercise

ECGs from anesthetized mice were recorded during echocardiography (Figure 3A), and the P, PR, QRS, QT, and RR intervals were measured and normalized to baseline (Figure 3B). Although we did not observe any significant changes in any of these parameters, a trend towards increased PR intervals was observed in male exercised mice alone (p=o.05). In female exercised mice, the QT interval displayed a shortening trend (p=0.05).

**Figure 3.**
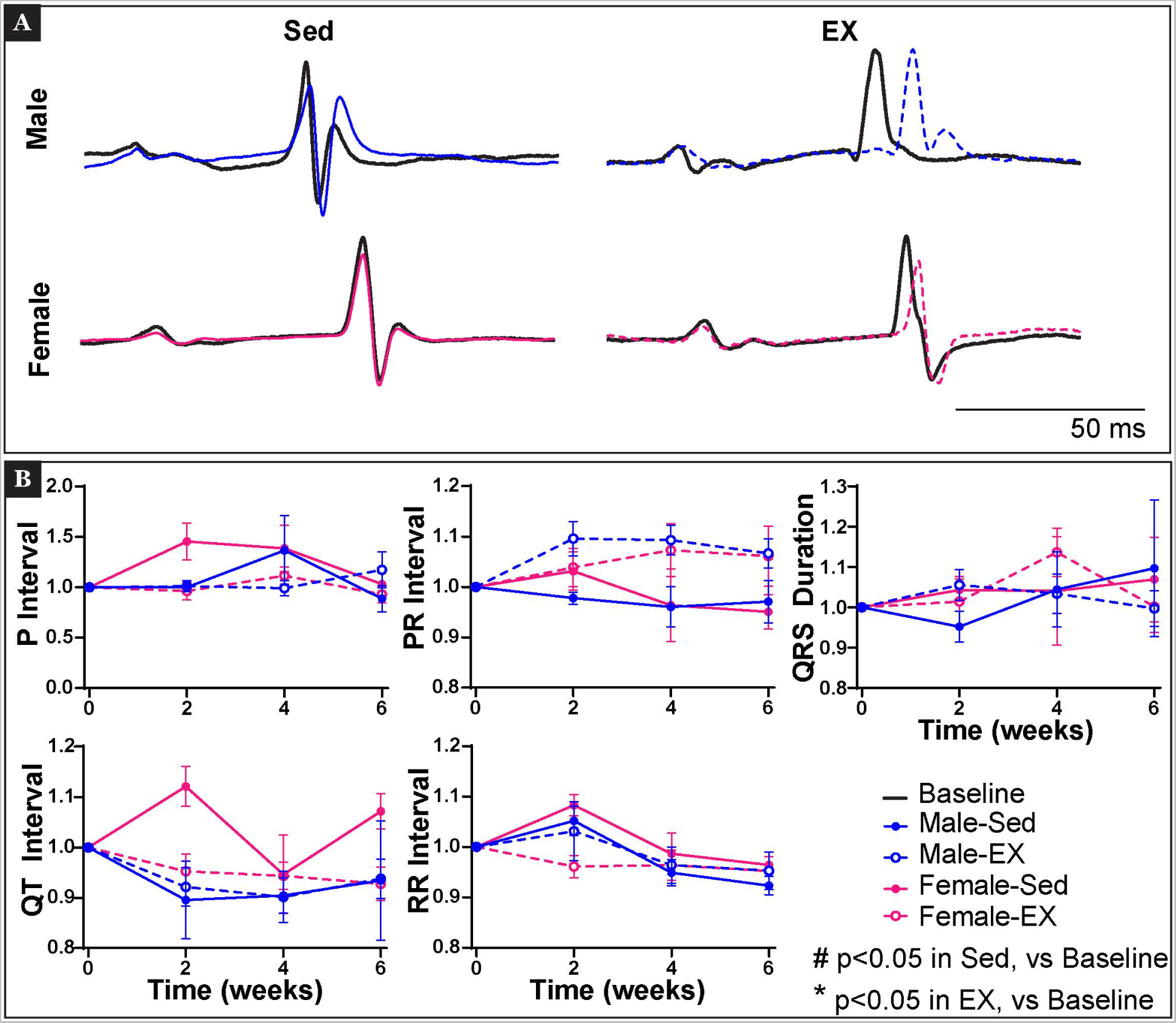
Electrocardiographic assessment of cardiac response to exercise. **A)** Electrocardiography was performed biweekly and representative ECGs from the male and female, Sed and EX mice at baseline (Week 0, black) and after the 6-week protocol (Week 6, blue: male, pink: female). Overlapping ECG traces are from the same mouse. **B)** Summary data of ECG parameters measured and normalized to each mouse’s own baseline value prior to start of experimental protocol is plotted over time. Three-way ANOVA tests were performed to determine significant differences in ECG data presented in panel B, with time, sex, and treatment as variables. Paired, two-tailed Student’s t-tests were performed for post hoc analysis to determine significant changes induced by experimental protocol in the same mouse over time and significant p values are indicated in the figure. Benjamini-Hochberg correction was applied for multiple comparisons correction. p<0.05 denotes significance. The sample size for each group is as follows: Male sedentary = 11, Male exercised = 9, Female sedentary = 11, Female exercised = 9.

Next, triple-parametric optical mapping was performed to investigate further the effects of exercise on cardiac electrical, calcium handling, and metabolic functions. NADH intensity maps, voltage, and calcium activation maps were generated from all four experimental groups (Figure 4A). Representative voltage and calcium traces are illustrated in Figure 4B. Ten cardiac function parameters were measured, and the percent change in exercised hearts versus sedentary controls is reported in Figure 4C. Yet again, exercise-induced modulation of cardiac function was sex-dependent. Specifically, in male exercised mice, the Ca τ was decreased versus sedentary males (38±16 vs 30±3 ms). On the other hand, in female exercised mice, APD prolongation (60±2 vs 72±7 ms) and reduction in V_m_-Ca delay (−7±3 vs −2±2 ms) were observed versus sedentary female mice. These significant changes in electrophysiological parameters are associated with > ±20% change versus control (Figure 4C, radial graph). Lastly, comparing exercise-induced modulation in male versus female mice, only the V_m_-Ca delay was significantly different.

**Figure 4.**
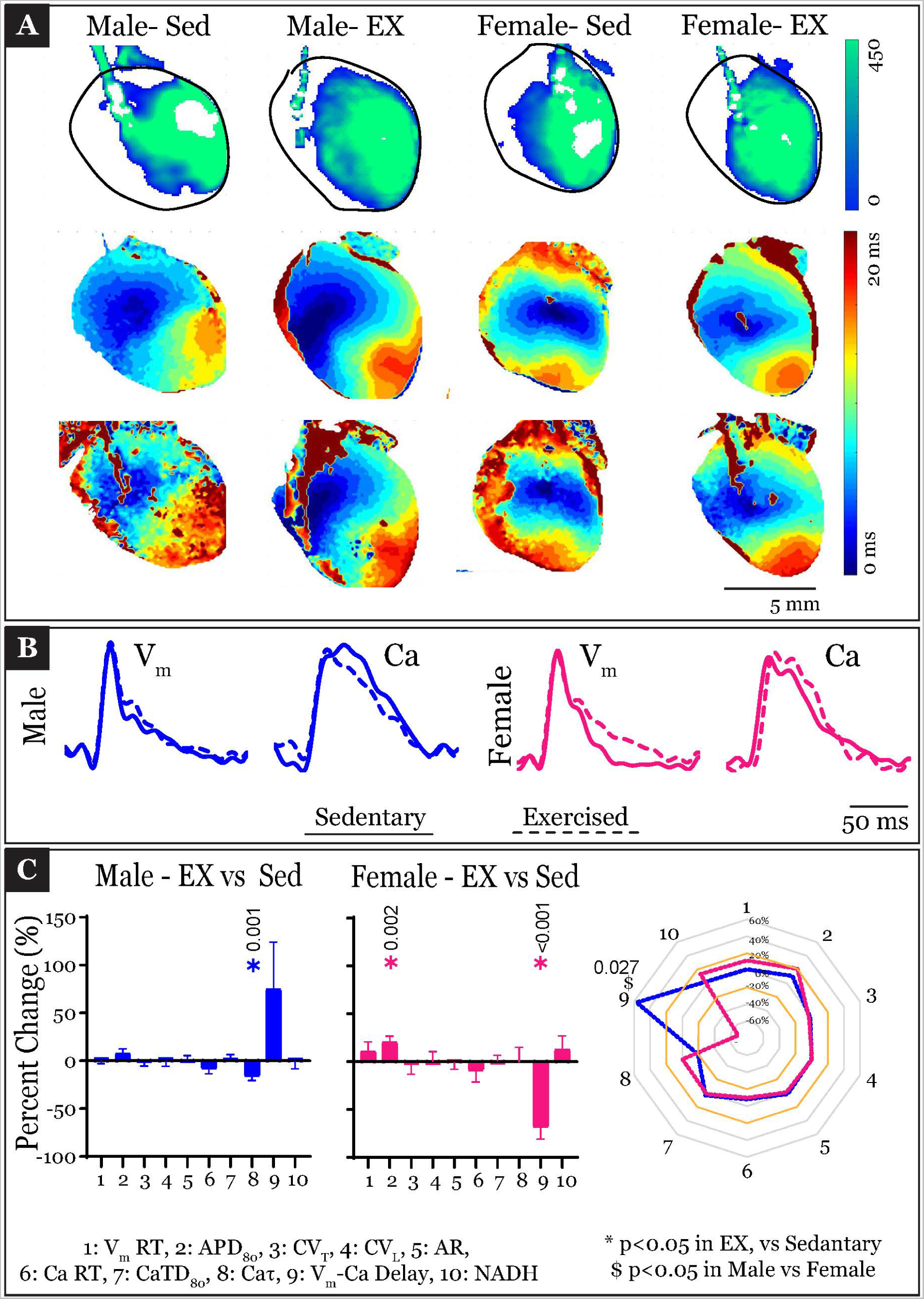
Cardiac electrical, calcium handling, and metabolic response to exercise. **A)** At the end of the experimental protocol, hearts were explanted, and triple parametric optical mapping was performed. Simultaneously recorded NADH intensity maps and voltage and calcium activation maps generated from triple-parametric optical mapping of male and female, Sed and EX hearts paced at 150 ms BCL are illustrated. All three maps in a given column are simultaneous recordings of the same field of view. **B)** Representative voltage and calcium optical traces from the optical mapping of hearts in the four experimental groups. Sedentary: solid line; exercised: dashed lines. **C)** Transmembrane potential, calcium handling, and metabolism-related parameters in exercised mice were normalized to sedentary controls and are illustrated as ten parameter panels and radial graph. Orange lines on the radial graph correspond with 20% and −20% change in any given parameter. V_m_ RT: rise time of the upstroke of action potential, APD_80_: action potential duration at 80% repolarization, CV_T_: transverse conduction velocity, CV_L_: longitudinal conduction velocity, AR: anisotropic ratio, CaRT: rise time of upstroke of calcium transient, CaTD_80_: calcium transient duration at 80% reuptake, Ca τ: decay constant of calcium transient, V_m_-Ca delay: delay between activation of voltage and calcium signals, NADH: normalized NADH intensity. Two-way ANOVA was performed to determine significant differences in data presented in Panel C, with sex and treatment Unpaired, two-tailed Student’s t-tests were performed to determine significant changes in optical mapping parameters presented in panel C and significant p-values are indicated. Benjamini-Hochberg correction was applied for multiple comparisons correction. p<0.05 denotes significance. The sample size for each group is as follows: Male sedentary = 5, Male exercised = 6, Female sedentary = 6, Female exercised = 6.

These data further support the observation that exercise-induced modulation of cardiac electrophysiology is sex-dependent.

### Exercise increased arrhythmia burden

Next, arrhythmia inducibility was tested in *ex vivo* hearts by programmed electrical stimulation. Representative traces illustrating sinus rhythm, VT, and VF are illustrated in Figure 5A. Burst pacing of exercised male and female hearts induced VT and VF, which was recorded by pseudo-ECGs (Figure 5B, left). VT and VF incidence and arrhythmia duration were quantified in each experimental group (Figure 5B, right), and phase maps were generated from optical recordings of the arrhythmia episodes (Figure 5C).

**Figure 5.**
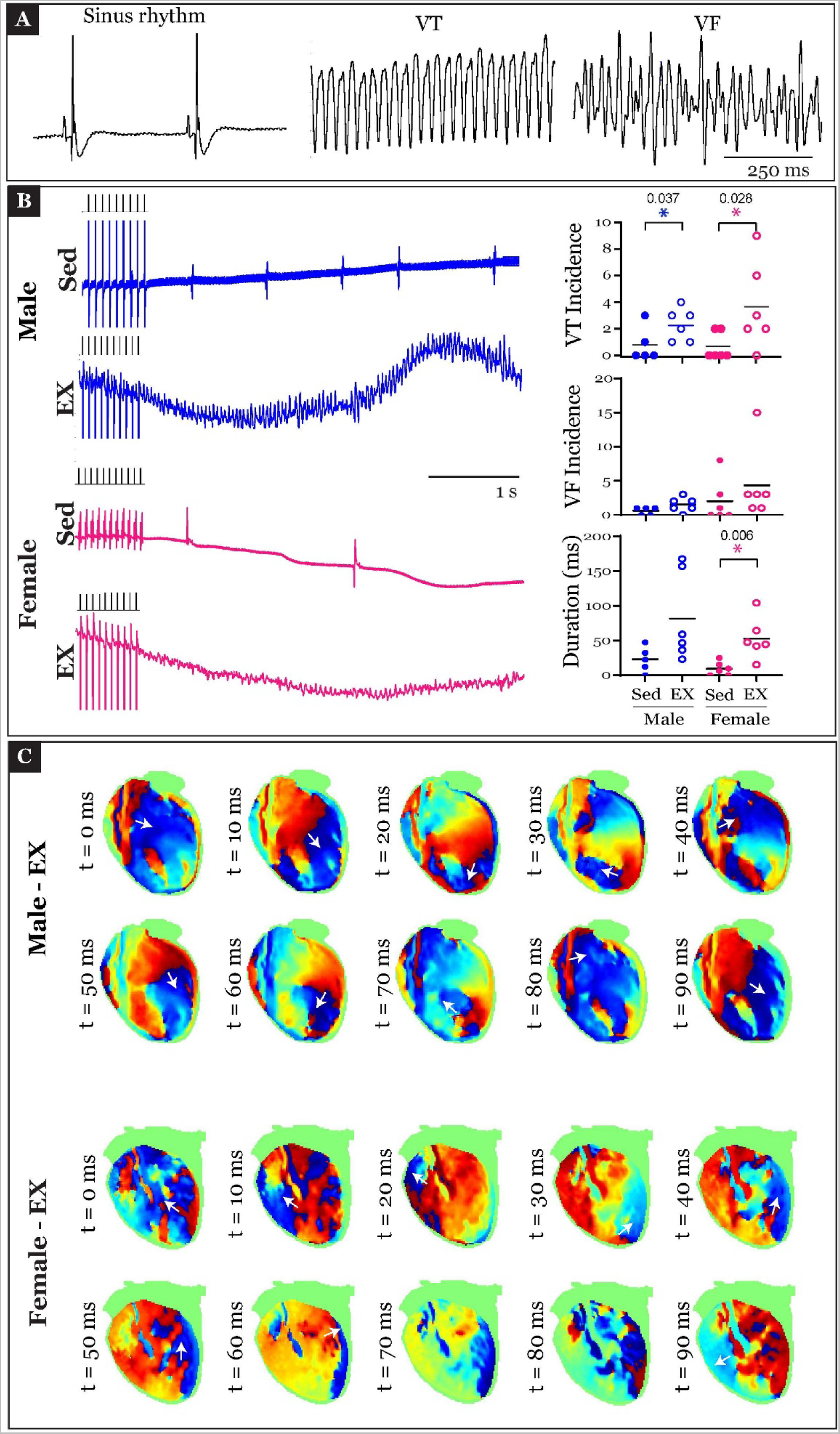
Increased arrhythmia susceptibility in hearts due to exercise. **A)** Representative ECG traces illustrating incidences of sinus rhythm, VT and VF in *ex vivo* mouse hearts. **B)** Pseudo-ECG traces illustrating burst pacing (indicated by black pulses above ECG traces) followed by either an arrhythmia episode or sinus rhythm in the four experimental groups (left) is illustrated. Quantification of VT and VF incidences and total arrhythmia duration in these four groups is plotted on the right. **C)** Phase maps were generated from optical recordings of arrhythmia episodes in exercised male and female *ex vivo* hearts. These maps, generated 10 s apart illustrate the propagation of the excitation wavefront (red regions) during the arrhythmia. White arrows indicate direction of propagation of excitation wavefront. In the male heart, a clear circular movement of the wavefront is evident indicating reentrant arrhythmia, whereas in the representative female heart, a more complex arrhythmia pattern is observed. Two-way ANOVA was performed to determine differences in data presented in Panel B, with sex and treatment as the two variables. Unpaired, two-tailed Student’s t-tests were performed as post hoc analysis, to determine significant changes in arrhythmia parameters presented in panel B. Benjamini-Hochberg correction was applied for multiple comparisons correction. p<0.05 denotes significance. The sample size for each group is as follows: Male sedentary = 5, Male exercised = 6, Female sedentary = 6, Female exercised = 6.

While VT incidences in exercised male and female hearts were significantly increased versus sedentary controls, VF incidences were not statistically different. Additionally, arrhythmia duration was also prolonged in exercised mice but was statistically significant only in female exercised mice compared to sedentary female mice (males: p=0.07, female: p=0.01). Lastly, arrhythmias observed in male exercised mice were associated with stable rotors, as observed from the phase maps, whereas arrhythmias in female exercised mice displayed a random activation pattern (Figure 5C).

### Sex-dependent gene expression modulation in response to exercise

Lastly, the molecular pathways underlying the sexually dimorphic modulation of cardiac physiological parameters were assessed at the transcriptional level. Not much is known about the changes in cardiac gene expression in response to moderate endurance exercise, particularly, sex differences in this phenomenon. Here, we present changes in genes associated with the cardiac structure, mechanics, and electrophysiology that could contribute to the functional effects described above. RNA sequencing was performed and an overview of the genetic remodeling in exercised mice vs sedentary controls is presented in Figure 6 and genetic sex differences is presented in Figure 7.

**Figure 6.**
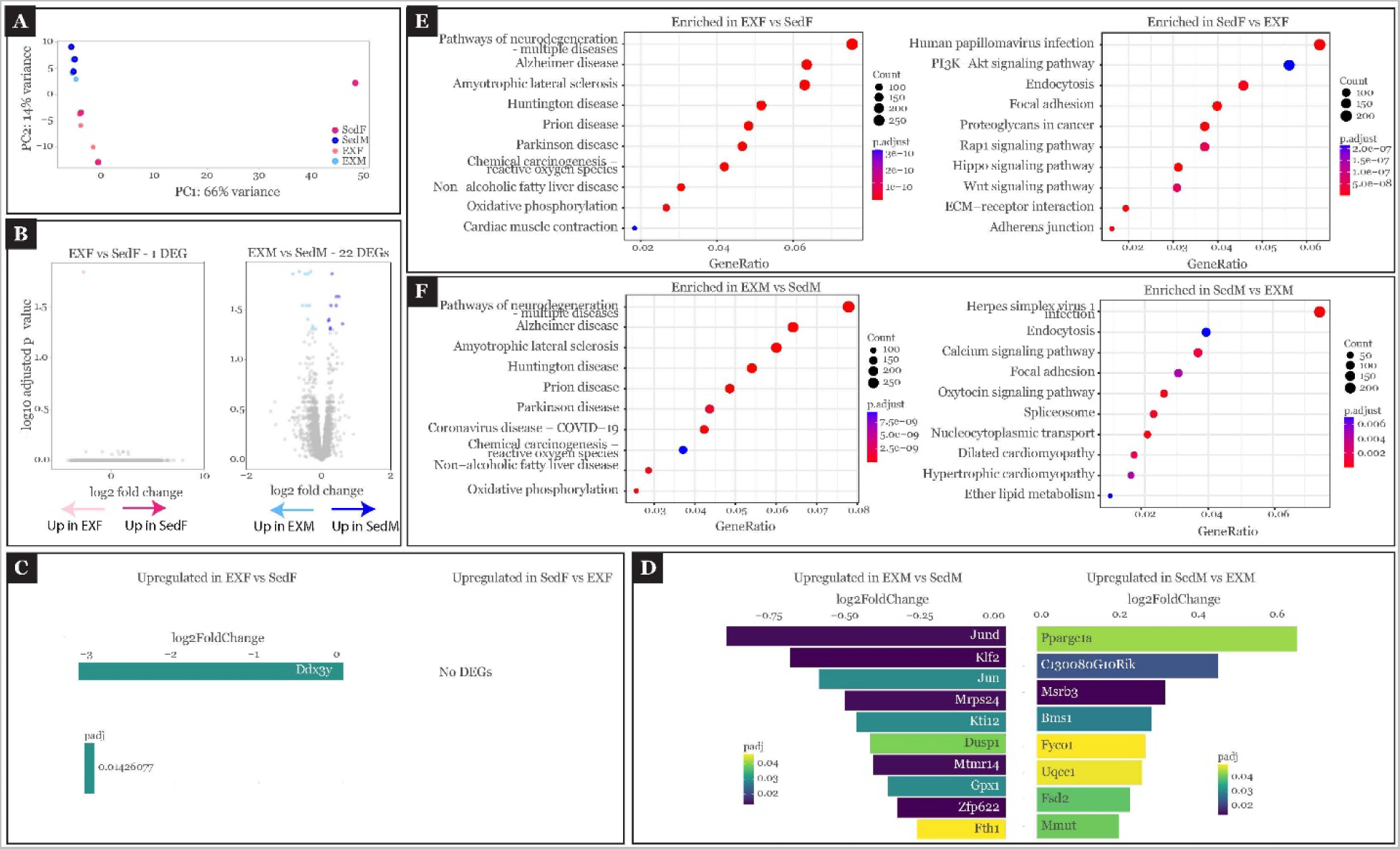
Differential cardiac gene expression in response to exercise. **A)** PCA plot illustrating the separation of sedentary and exercised groups in male and female gene expression profiles. **B)** Volcano plots showing DEGs in exercised versus sedentary controls. **C,D)** Top ten DEGs in exercised vs sedentary mice of either sex. In instances where fewer DEGs were identified between groups, only those genes are shown. **E,F)** Top enriched pathways in exercised vs sedentary mice of either sex. DESeq2 package in R was used to determine significant DEGs. clusterProfiler package in R was used to determine enriched pathways. The significance level was set to < 0.05 and log2 fold change of > 0 was reported as upregulated and < 0 as downregulated. The sample size for each group is as follows: Male sedentary = 3, Male exercised = 3, Female sedentary = 3, Female exercised = 3.

**Figure 7.**
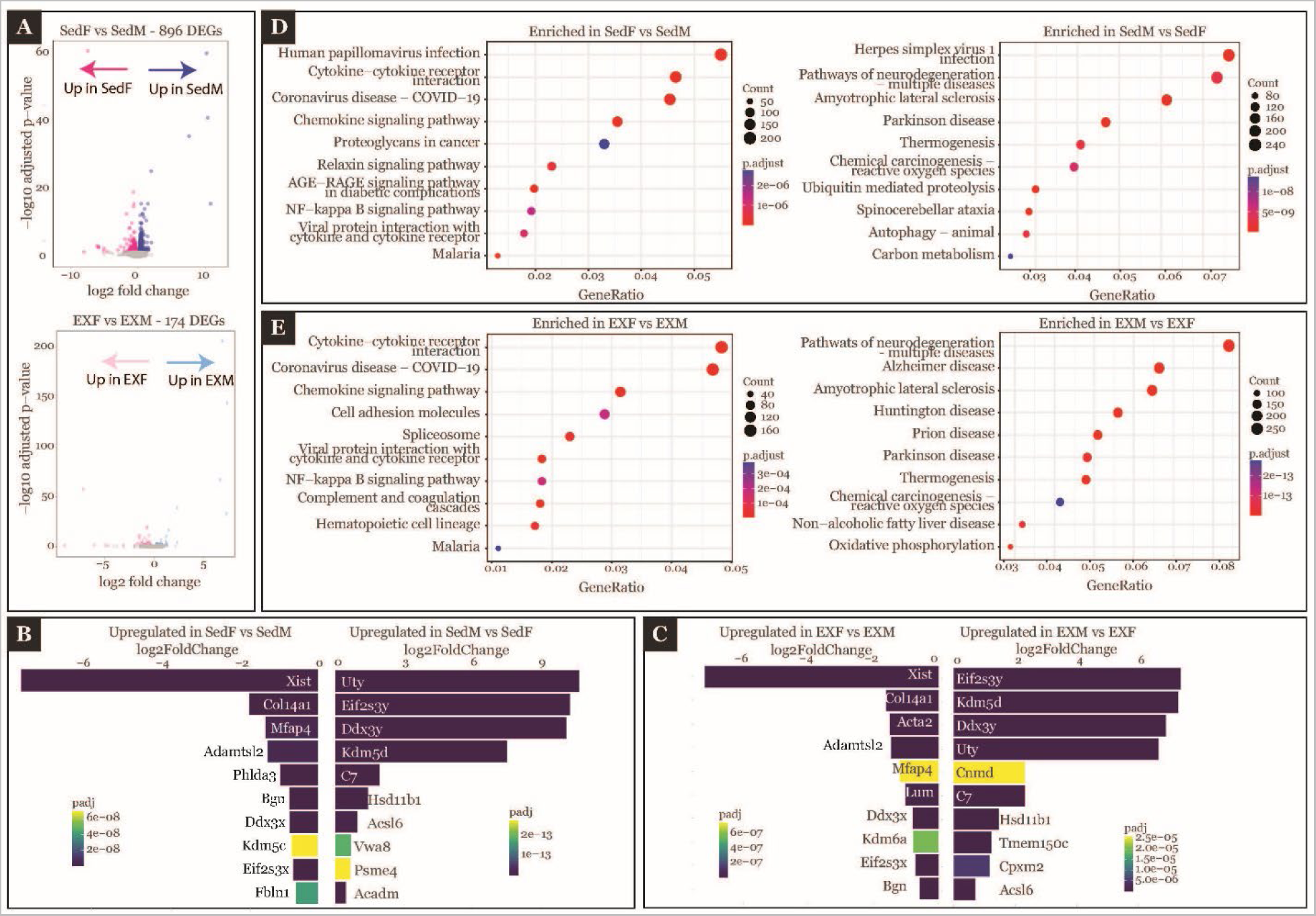
Sex differences in cardiac gene expression in exercised and sedentary mice. **A)** Volcano plots showing sex differences in DEGs exercised and sedentary mice. **B, C)** Top ten DEGs in female vs male mice of exercised and sedentary groups. **D, E)** Top enriched pathways in female vs male mice of exercised and sedentary groups. DESeq2 package in R was used to determine significant DEGs. clusterProfiler package in R was used to determine enriched pathways. The significance level was set to < 0.05 and log2 fold change of > 0 was reported as upregulated and < 0 as downregulated. The sample size for each group is as follows: Male sedentary = 3, Male exercised = 3, Female sedentary = 3, Female exercised = 3.

The principal component analysis (PCA) plot demonstrates separation of male and female hearts along the first and second principal components but not much differences between exercised mice and sedentary controls of either sex (Figure 6A). Volcano plots in Figures 6B also illustrate fewer significant DEGs between exercised mice and sedentary controls (Males: 22 DEGs, Females: 1 DEG). Ddx3y gene was the only significantly upregulated gene in exercised female mice compared to sedentary female mice (Figure 6C). Interestingly, ddx3y is a Y-linked gene and should not be expressed in female mice. However, previous studies have reported the presence of ddx3y in female mice tissue due to sequence similarity of this gene with other autosomal genes and the cross-reactivity of the ddx3y probe with these other genes. (27) Comparing exercised male mice to sedentary males (Figure 6D), significant DEGs included GPX1, Mtmr14 and Mrps24 which are involved in metabolic processes as well as Klf2 and Fth1 which are involved in immune responses. KEGG analysis revealed several cellular processes that were modulated by exercise (Figure 6E and 6F). Oxidative phosphorylation related genes were enriched in both male and female exercised mice compared to sedentary controls. Additionally, in female exercised mice, genes related to cardiac muscle contraction were also enriched.

Finally, comparing females to males in each group (exercise and sedentary) revealed a larger number of significant DEGs (Sedentary: 896 DEGs, Exercised: 174 DEGs, Figure 7A). In female mice, Ddx3x gene which is involved in immune response was upregulated in both sedentary and exercised groups. (Figure 7B, C). Additionally, Acta2 gene which is involved in muscle contraction is also upregulated in exercised female mice vs exercised males. (Figure 7C). In male mice, lipid metabolism related genes Hsd11b1 and Acsl6 were among the top DEGs in both sedentary and exercised groups. KEGG analysis (Figure 7D and 7E) revealed enrichment of genes involved in infectious diseases and immune responses in female mice regardless of exercise or sedentary group. On the other hand, in male mice, enriched genes were associated with neurological diseases such as Parkinson, Huntington and Alzheimer’s disease. Genes related to oxidative phosphorylation were also enriched in exercised male mice compared to exercised female mice.

KEGG pathway maps of top modulated pathways and a full list of top DEGs are included in Supplements 1 and 2, respectively.

## DISCUSSION

This study, for the first time, demonstrates that moderate-intensity endurance exercise modulates multiple aspects of cardiac anatomy and physiology in sex-specific manner (Figure 8). Briefly, the effects of exercise were predominantly observed in female mice and not in males. Exercise caused LV dilatation, changes in cardiac mechanical function, and arrhythmogenic remodeling of cardiac electrophysiology. This resulted in increased duration and complexity of arrhythmias in female mice. These functional modulations were associated with the genetic remodeling of genes related to skeletal muscle, contraction, and metabolism. Together, this data highlights the beneficial and detrimental effects of exercise on male and female cardiac physiology.

**Figure 8.**
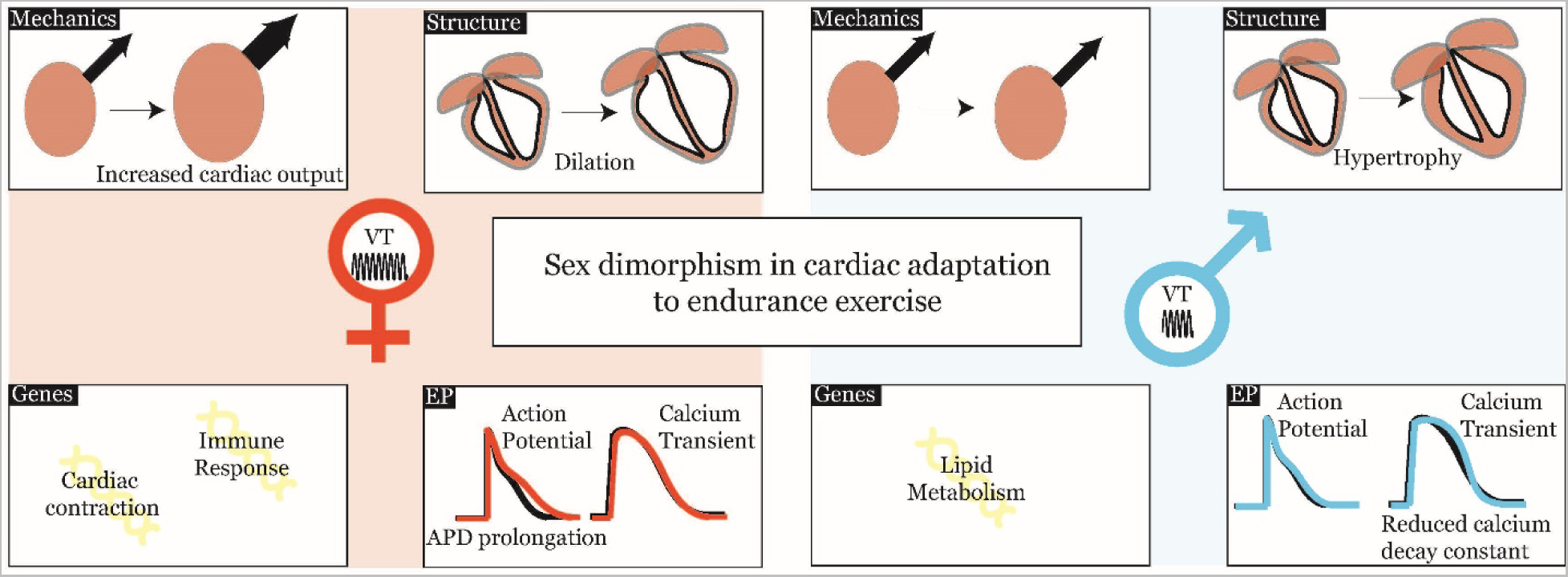
Summary of Results. Illustration of major findings of the study in female (left, red background) and male (right, blue background) mice.

### Structural remodeling

High-intensity and prolonged exercise in athletes has been reported to cause mostly reversible, physiological remodeling of cardiac structure. This includes increased fibrosis, enlargement of the LV chamber (dilatation), hypertrophy, increased mass etc.(5,14) The increased demand on the heart is an underlying cause for these effects and this has been reported as early as 1918 by Thomas Lewis based on his study of remodeling in soldiers’ hearts. He refers to this condition as the effort syndrome.(28) However, little is known about the effects of moderate-intensity exercise on non-athletes. Here, we report that moderate endurance exercise can also cause remodeling of the cardiac chambers, particularly the LV. Echocardiographic imaging of the LV identified different responses in male and female hearts to exercise. While LV dilatation was observed in female hearts, LV hypertrophy was indicated in male hearts. The female heart responded to exercise with an enlarged LV cavity and thinning of the LV wall. The enlarged cavity, which suggests increased diastolic filling, could be a response to increased demand for blood flow during exercise. On the other hand, male hearts responded to exercise with increased LV wall thickness.

These structural changes correlated with the genetic remodeling observed in the exercised mice. Acta2 gene associated with actin myofilament structure was among the top DEGs in female exercised mice, suggesting structural remodeling at the macroscopic and microscopic levels. It also indicates that the effects of exercise on female hearts could be both a response to functional demands and genetic remodeling. It would therefore be interesting to see the reversibility of these effects in future studies.

### Mechanical Remodeling

The relationship between exercise and cardiac mechanics is complex. On the one hand, impaired cardiac mechanics in patients with CVD can lower exercise capacity.(29) On the other hand, exercise can improve cardiac mechanics over time in patients with CVDs such as heart failure and coronary artery disease.(11,30) One commonly measured parameter of mechanical function, EF, is reduced in athletes.(31) However, what distinguishes the EF response in healthy athletes versus CVD patients with dilated cardiomyopathy is the ability of EF to increase during exercise in athletes.(32)

In this study, we measured EF at rest in exercised mice. The findings suggest that even moderate levels of endurance exercise can reduce EF. However, EF reduction was only observed in female mice. Reduction in EF in athletes does not necessarily signify mechanical impairment. EF is the ratio of the amount of blood ejected per beat (SV) to the total diastolic volume. Based on the LV dilatation observed in the exercised mice, it can be stated that diastolic volume is increased in female exercised mice. Our findings also identified that SV remains stable after 6 weeks of exercise. Therefore, the reduction in EF is possibly due to the enhanced diastolic volume and not due to reduced ejection volume per beat.

We also report an increase in CO in female exercised mice. CO is the product of SV and HR. In our results, we report non-significant increases in both SV and HR after week 2, which resulted in increased CO at weeks 4 and 6 in female exercised mice. This contradicts previous reports of decreased resting HR and CO in athletes.(33) Several factors could contribute to this discrepancy, 1) differences between moderate and intense exercise in athletes, 2) mouse versus human response to exercise, 3) sex differences (previous studies did not consider the sex of the athlete), and 4) short duration (6 weeks) of exercise regimen in the current study. Additionally, the HR reported here was recorded from anesthetized mice, and we have previously demonstrated that anesthesia can modulate response to treatments in mice.(24)

Looking at the whole picture of cardiac mechanical function, moderate endurance exercise in mice appears to mimic some effects of extreme exercise, such as reduced EF, while having some opposing responses, such as increased HR. Regardless, the critical point here is that the heart adapts to the demands of moderate endurance exercise by increasing blood supply (increased CO). Importantly, this adaptation was only observed in female mice and not males after 6 weeks of exercise training.

Lastly, these changes in mechanical function correlated with changes in the expression of genes associated with cardiac muscle contraction, as well as actin myofilament structure. These genes were enriched in female exercised hearts versus sedentary controls, but not in males.

### Electrical Remodeling

ECGs from athletes have a different set of evaluation criteria because of the significant electrical remodeling that occurs in response to intense exercise.(34,35) These include sinus bradycardia, 1^st^ and 2^nd^ degree AV block, T wave inversion, high voltage QRS, and others. In this study, we aimed to identify the effects of moderate endurance exercise on cardiac electrophysiology. A trend in increasing PR interval was observed in male exercised mice alone which could indicate AV node remodeling. However, this remodeling was not as drastic as to cause AV block, as is the case with high-intensity exercise in athletes. Furthermore, it needs to be noted that no changes in PR interval were observed in female exercised mice. This sex difference in the effects of exercise on AV node function agrees with a previous finding that AV block was less prevalent among female Olympic athletes.(34) This finding could have significant implications in diagnosing electrical remodeling in men versus women athletes and non-athletes who undergo moderate endurance exercise.

ECG analysis gives an understanding of exercise-induced global changes in cardiac electrophysiology, while assessment on a microscopic scale will provide more evidence on the mechanisms underlying these global changes. Therefore, we further examined ventricular electrical, calcium handling, and metabolic functions by triple-parametric optical mapping. Regarding electrical function, we observed APD prolongation in *ex vivo* female exercised mice hearts. This is an interesting finding because previous studies have reported QT interval shortening in response to exercise, and this current study also displays a trend of QT shortening (p=0.05 at Week 6) in female exercised mice. QT shortening in athletes is a response to an increased vagal tone.(36) However, when vagal stimulation is removed, as in the explanted Langendorff perfused heart for optical mapping, we observed an increase in APD. This suggests that exercise has dual and opposing effects on APD. On the one hand, ion channel remodeling could underlie APD prolongation, while increased vagal tone could promote APD shortening. This could have severe implications for arrhythmogenicity in female exercised hearts. Regarding calcium handling, the Ca τ was reduced in male exercised mice, which indicates that the reuptake of calcium back into the sarcoplasmic reticulum or the removal of the calcium ions from the cytoplasm by the sodium-calcium exchanger is faster. This could provide some protection to the male exercised hearts by preventing cytosolic calcium buildup during faster heart rates and could underlie shorter arrhythmia incidences in these male exercised mice. While previous studies have reported that testosterone increases sarcoplasmic/endoplasmic reticulum calcium ATPase 2a (SERCA2A) and thereby faster calcium reuptake in males,(37) the effects of exercise on this phenomenon in males require further investigation. Regarding metabolic function, no significant changes were observed in total NADH autofluorescence intensity in either male or female exercised mice versus sedentary controls. NADH accumulation in the heart occurs when the metabolic respiration rate decreases under conditions such as ischemia, while NADH reduction is observed in conditions of high metabolic respiration rates, such as during exercise. While chronic exercise training could lead to significant metabolic adaptations to respond to bouts of exercise, our results indicate that metabolic states were similar between our sedentary and exercised groups in resting state in these explanted mechanically unloaded hearts. However, the significant remodeling of metabolism-related genes, particularly in male exercised hearts, suggests that this aspect needs further investigation.

Probably due to electrical and calcium handling functions remodeling, both male and female exercised mice hearts were more susceptible to arrhythmias. VT was the most prevalent form of arrhythmia in these hearts, while some episodes of VF were also observed. Furthermore, the duration and complexity of arrhythmia patterns were higher in female exercised hearts.

### Sex dependence of exercise-induced remodeling

Throughout the discussion of the results of this study, one common factor has surfaced as an important modulator of the effects of exercise – sex differences. To our knowledge, this is the first study to report sex differences in the modulation of cardiac physiology and transcriptome in response to moderate levels of endurance exercise. The question that then arises is what underlies these sex differences.

It is well-established that sex hormones play a prominent role in modulating cardiac function.(38,39) Both estrogen and testosterone modulate cardiac ion channels function through genomic and non-genomic mechanisms.(40) Estrogen reduces cardiac potassium currents such as the transient outward potassium current (I_to_), rapid and slow delayed rectifier potassium currents (I_Kr_ and I_Ks_, respectively) which could produce APD prolongation.(41–44) Contrarily, testosterone increases I_Kr_, I_Ks_ and the inward rectifier potassium current (I_K1_), resulting in APD shortening.(45) Testosterone also reduces the L-type calcium current (I_CaL_) which could further shorten APD while the effects of estrogen on I_CaL_ is cardiac region-specific.(46,47) Finally, testosterone has also been reported to increase rates of contraction and relaxation which could be attributed to increased ryanodine receptors 2 (RyR2) and SERCA2A.(37)

Exercise modulates estrogen and testosterone levels depending in the type, intensity, duration and timing after exercise. Endurance exercise decreases estrogen in women while it remains unaltered in men.(48,49) On the other hand, endurance exercise acutely increases testosterone in men and women but chronically, reduces testosterone compared to sedentary subjects.(50,51) While the effects of sex hormones on cardiac ion channels are well-established, their effects in endurance exercise trained subjects are not known. In this study, we report that exercise produced sex-dependent modulation of cardiac electrophysiology and arrhythmias. While arrhythmia incidence in male and female mice were increased by exercise training, female mice had arrhythmias of longer duration. The role of sex hormones in modulating cardiac electrophysiology in exercise trained subjects requires further investigation.

Another crucial underlying mechanism for sex differences in cardiac physiology modulation by exercise could be cardiac metabolic differences. Metabolism of fatty acids and glucose produces ATP required for the functioning of contractile machinery, ion channels, and pumps. Sex differences in cardiac metabolism exist in healthy adults, where women depend more on fatty acids for metabolism than men.(16) Exercise increases metabolic rate to keep up with the increased demands of the heart, and more importantly, it further amplifies the sex differences in cardiac metabolism. Thus, sex differences in cardiac metabolism could also contribute to the results reported in this study, which will be the focus of future studies.

## Acknowledgements

None.

**Supplement1.pdf: KEGG Maps.** KEGG analysis pathway maps for key modulated pathways such as adrenergic signaling, cardiac muscle contraction, oxidative phosphorylation, calcium signaling, PI3K-AKT signaling, and WNT signaling in sedentary and exercised, male and female mice.

**Supplement2.xls: List of top DEGs.** Gene names and gene ontology terms associated with all genes listed in figures 6 and 7.

**Supplement 3: Statistical Analysis.** Numerical values of data presented in figures presented as mean ± standard deviation, and results of two- and three-way ANOVA tests performed on these data.

