## Supplementary figures and images for "Moderate Endurance Exercise Increases Arrhythmia Susceptibility and modulates Cardiac Structure and Function in a Sexually Dimorphic manner"

### Supplement 1

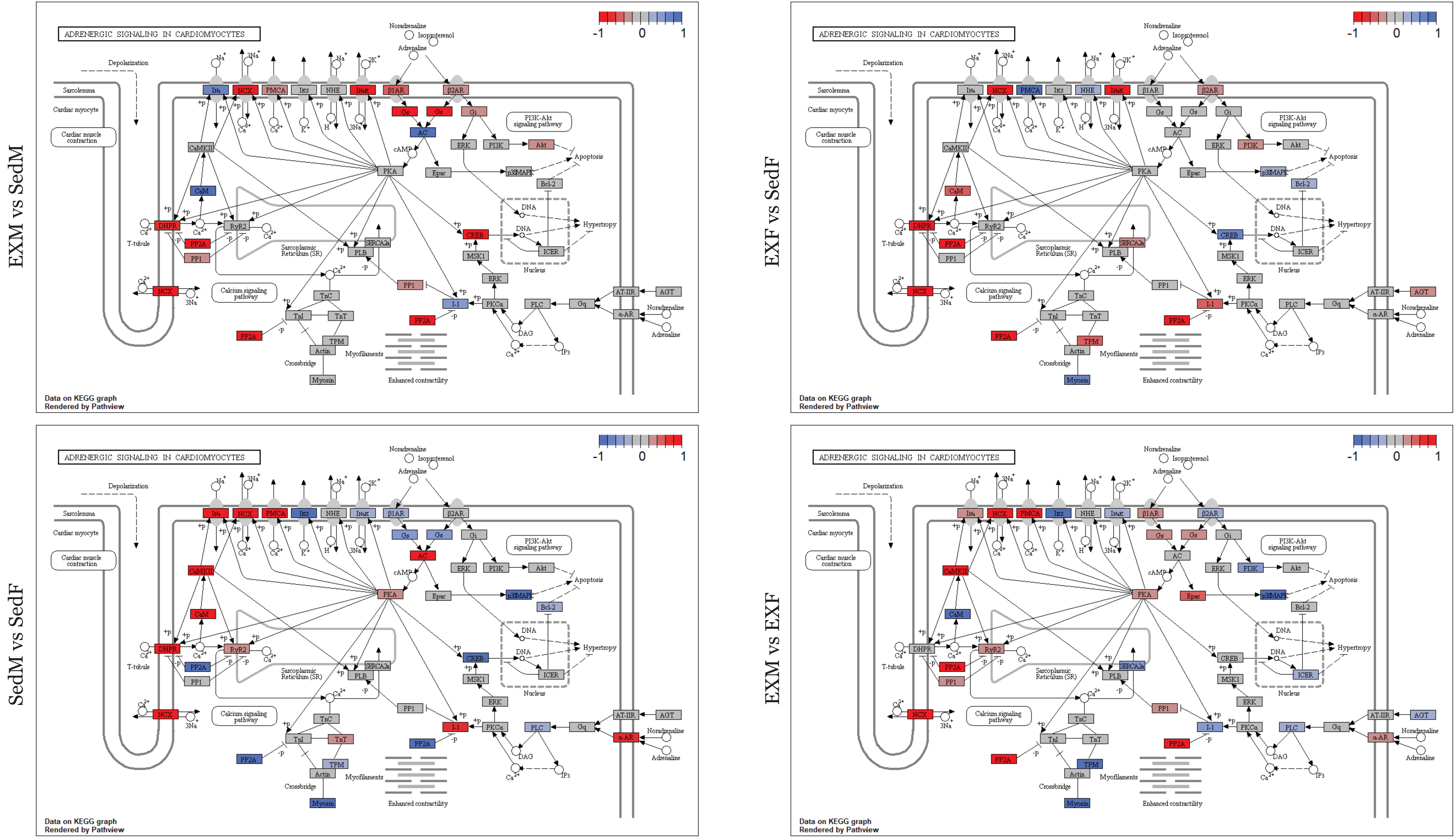

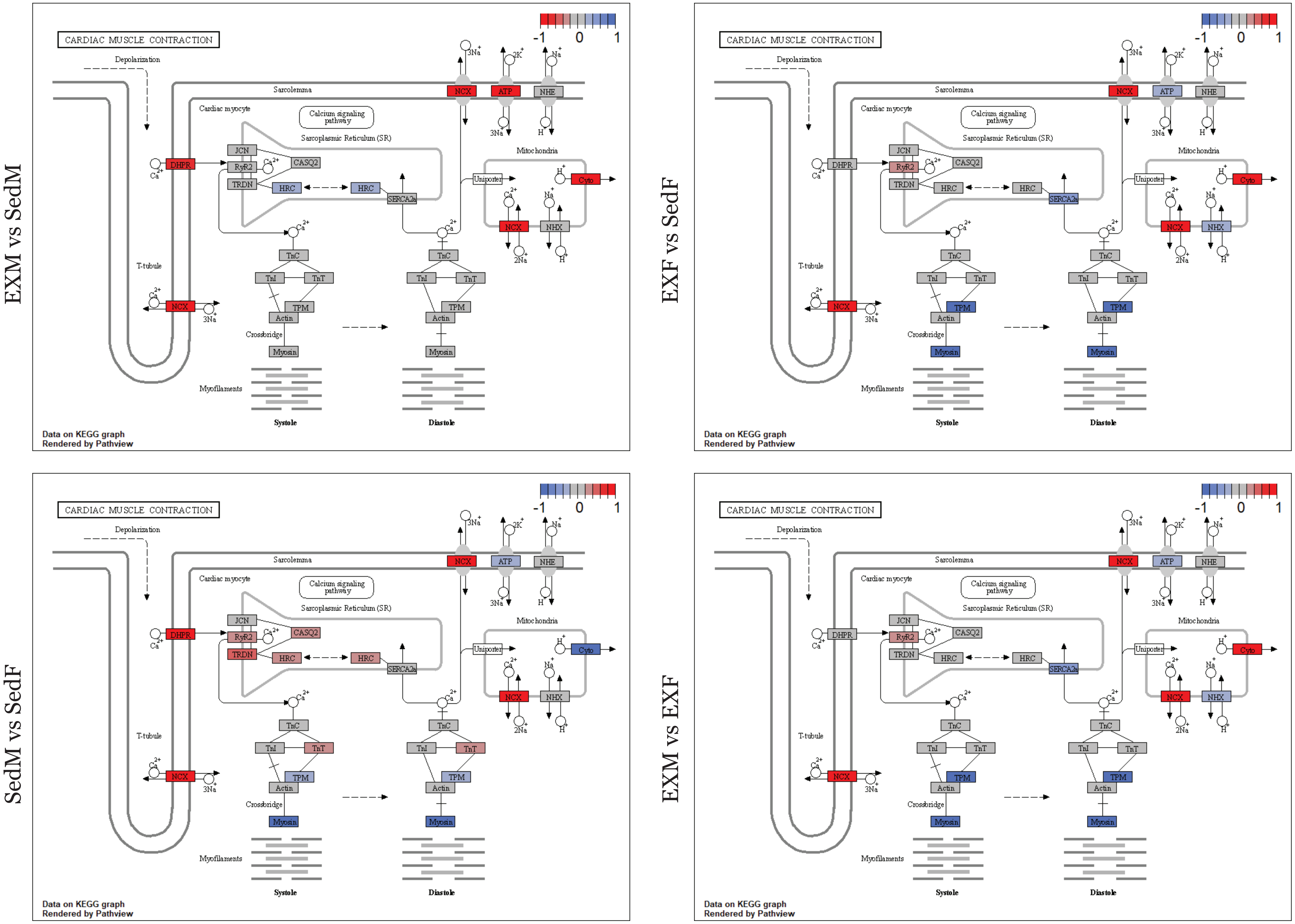

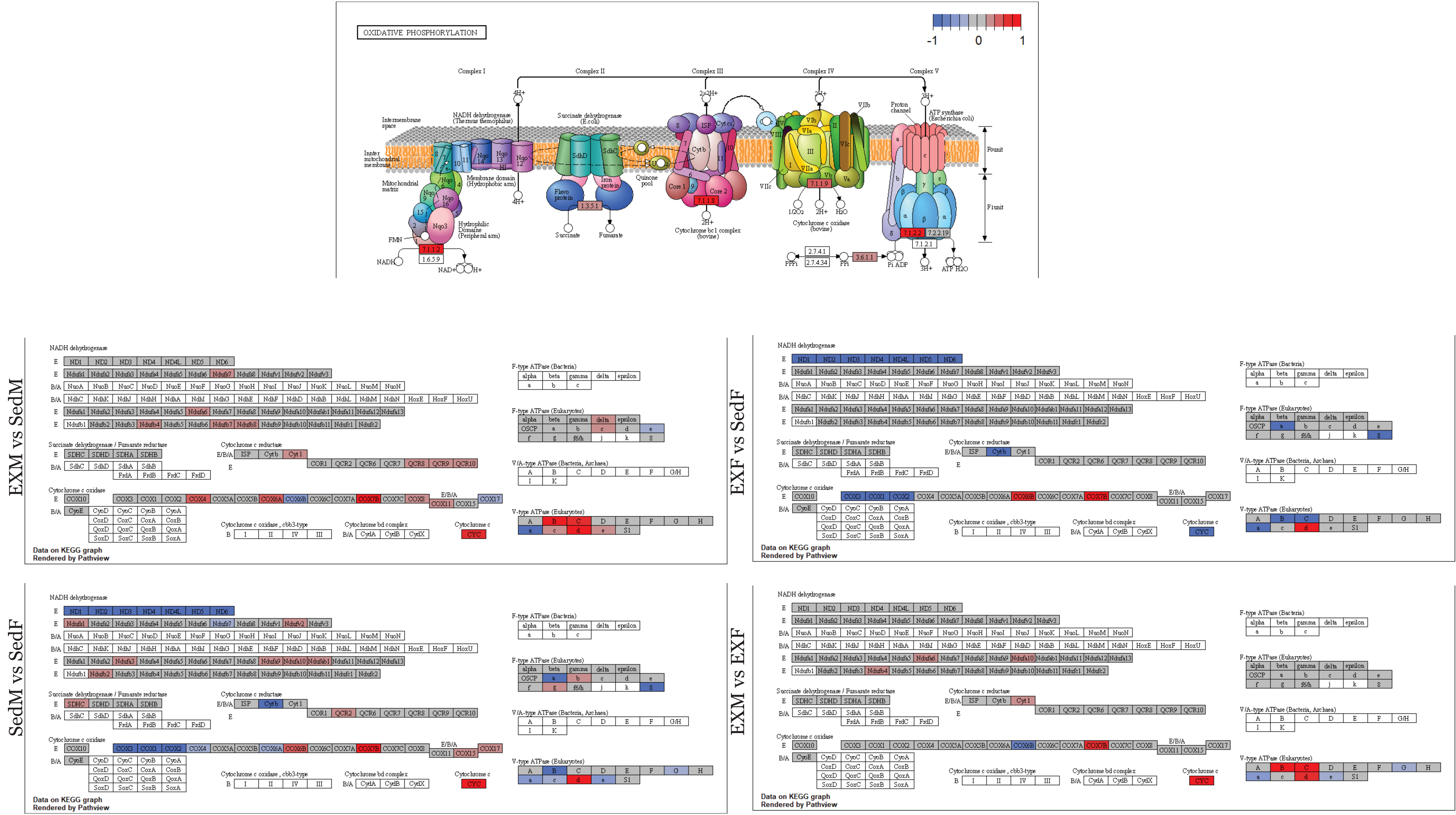

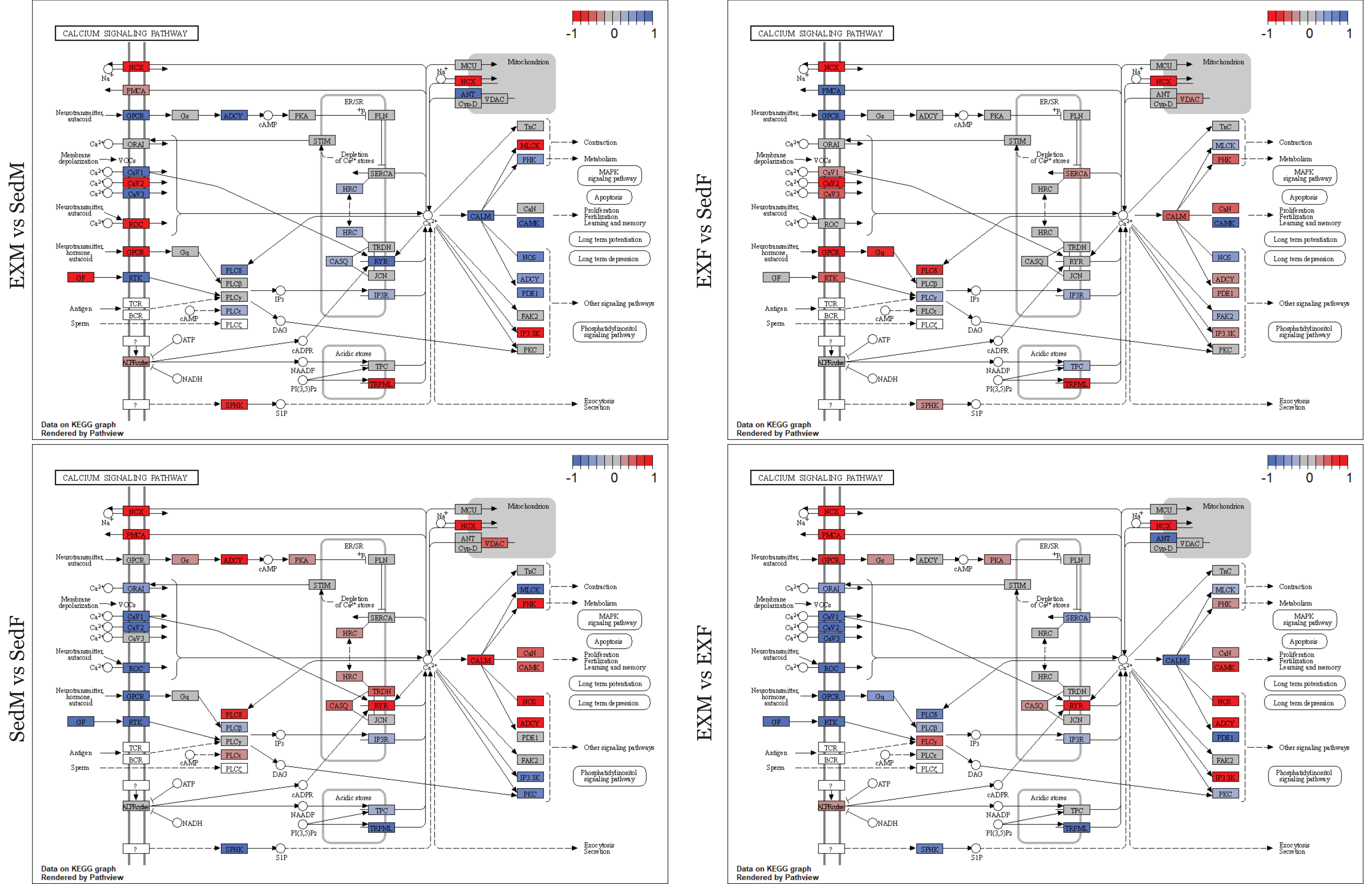

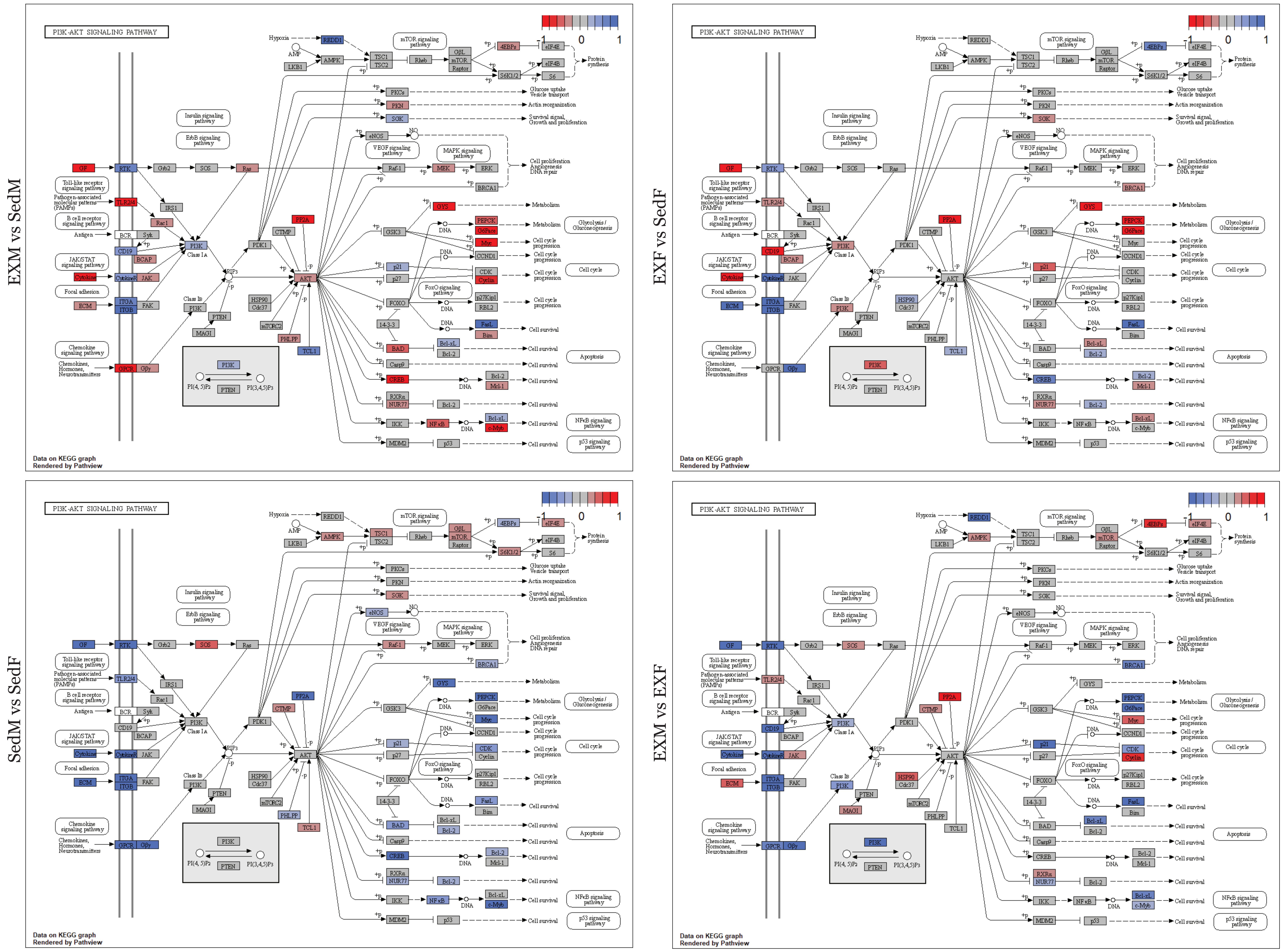

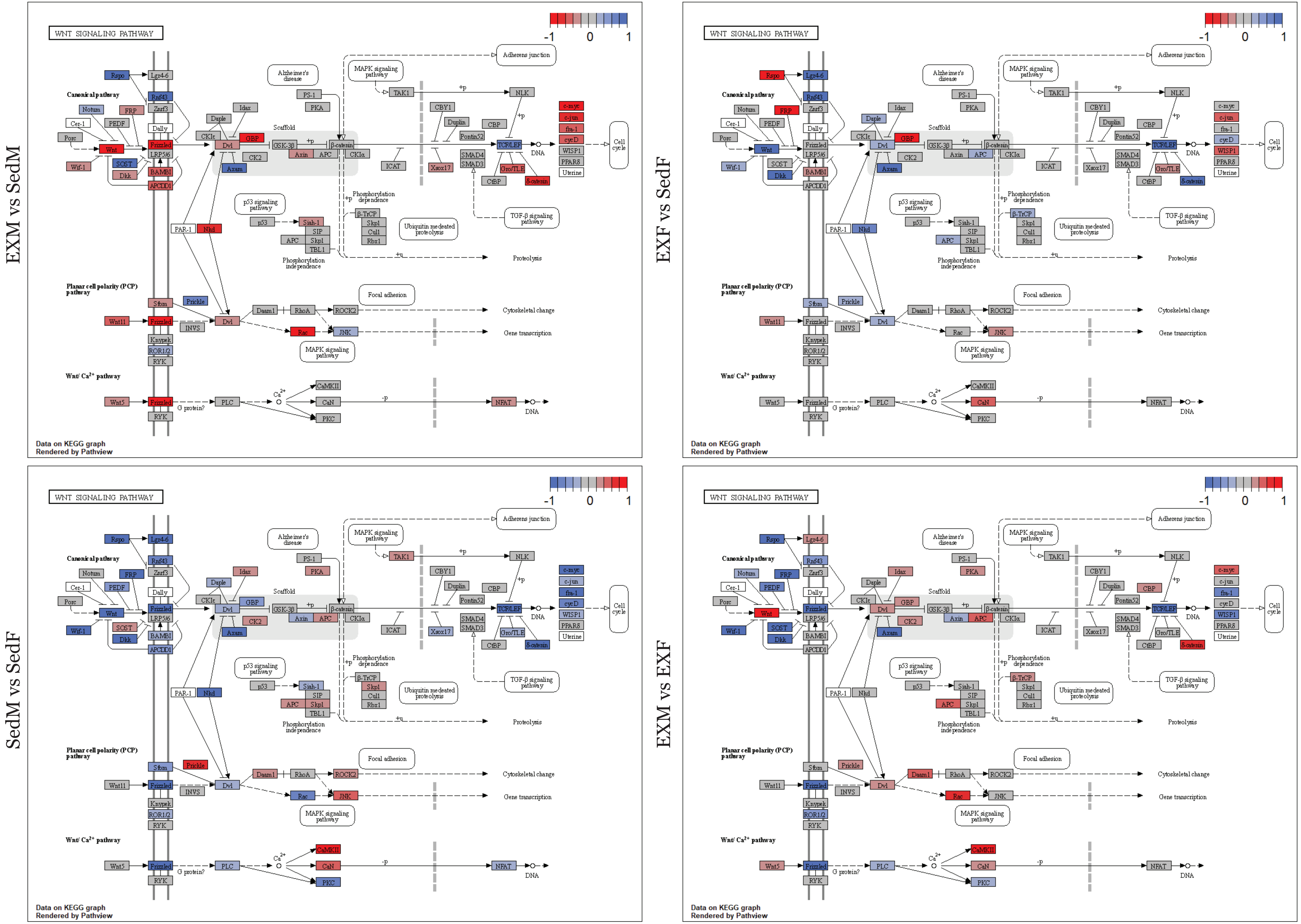
